# Design, development, and evaluation of gene therapeutics specific to KSHV-associated diseases

**DOI:** 10.1101/2025.02.19.639178

**Authors:** Tomoki Inagaki, Jonna Espera, Kang-Hsin Wang, Somayeh Komaki, Sonali Nair, Ryan R. Davis, Ashish Kumar, Ken-ichi Nakajima, Yoshihiro Izumiya

## Abstract

Kaposi’s sarcoma-associated herpesvirus (KSHV) is the causative agent of Kaposi’s sarcoma (KS) and two human lymphoproliferative diseases: primary effusion lymphoma and AIDS-related multicentric Castleman’s disease. KSHV-encoded latency-associated nuclear antigen (LANA) is expressed in KSHV-infected cancer cells and is responsible for maintaining viral genomes in infected cells. Thus, LANA is an attractive target for therapeutic intervention for KSHV-associated diseases.

Here, we devised a cancer gene therapy vector using the adeno-associated virus (AAV), which capitalizes the LANA’s function to maintain terminal repeat (TR) containing circular genome in latently infected cells and the TR’s enhancer function for KSHV inducible gene promoters. By including two TR copies with a lytic inducible gene promoter (TR2*-OriP*), we prepared an AAV vector, which expresses an engineered thymidine kinase (TK) selectively in KSHV-infected cells. Ganciclovir (GCV), an anti-herpesvirus drug, effectively eradicated multiple KSHV-infected cells that include iPSC-derived epithelial colony-forming cells, but not non-KSHV-infected counterparts in the presence of AAV8-TR2*-OriP*-TK. In addition, AAV8-TR2*-OriP*-TK prevents KSHV virion production from reactivated cells, spreading KSHV infections from reactivated cells. Anti-cancer drugs, known to reactivate KSHV, stimulated TK expression from the vector and, therefore, synergized with AAV8 TR2*-OriP*-TK to induce KSHV-infected cancer cell death. Finally, the AAV8-TR2*-OriP*-TK with GCV completely diminished KSHV-infected cancer cells in the xenograft tumor model. The new cancer gene therapeutics should augment the current clinical protocol for KS.

## Introduction

Kaposi’s sarcoma-associated herpesvirus (KSHV) was discovered in 1994 and is one of the eight human herpesviruses. KSHV is the causative agent of Kaposi’s sarcoma ^1, 2^, two human lymphoproliferative diseases, primary effusion lymphoma (PEL) ^3, 4^, AIDS-related multicentric Castleman’s disease (MCD) ^5, 6^, and a more recently described interleukin-6 related disease, KSHV-inflammatory cytokine syndrome (KICS) ^7, 8^. These highly inflammatory diseases are a leading cause of cancer deaths in AIDS patients in sub-Saharan Africa. KSHV-encoded Latency-associated nuclear antigen (LANA) is frequently identified in KSHV-infected tumor cells. Also, KSHV LANA plays a role in KSHV-mediated tumorigenesis by manipulating cell cycle machinery ^9^. Accordingly, we focus on the LANA protein as a therapeutic target to inhibit tumorigenesis and KSHV replication ^10, 11^. However, we have yet to develop effective small-molecule inhibitors for LANA; we need to explore additional directions.

The KSHV viral genome consists of an approximately 140 kb unique coding region flanked by large copies of high G+C 801 bp terminal repeat (TR) ^12^. KSHV genomes persist in latently infected cells as circular genomes (episome) via tethering to the host cell chromosomes ^13^. During latency, a few latent genes are actively transcribed ^14^. Among these latent genes, ORF73 encodes LANA, which plays a crucial role in latent episome replication and maintaining the episome in daughter cells. The TR contains a DNA replication origin, which consists of two LANA-biding sites (LBS): a higher affinity site (LBS1) and a lower affinity site (LBS2) followed by an adjacent 32-bp GC-rich segment ^14^. Episome maintenance requires at least two copies of TR (LBS1/2 binding sites) ^12, 13^. DNA binding induces oligomerization of LANA_DBD_, and a hydrophobic interface between LANA dimer forms the decametric ring and is essential for cooperative DNA binding, hence episome maintenance ^15^. These mechanisms substantially increase LANA concentration around the TR, and the TR/LANA complexes can be seen as LANA dots in KSHV-infected cells with immunostaining^16^. The LANA dots (also called LANA nuclear bodies) are used to diagnose KSHV etiology ^17^.

In addition to being a LANA binding sequence, KSHV TR was found to function as a gene enhancer for inducible viral gene promoters ^18, 19^. Gene enhancers are a crucial regulatory genomic domain for differential gene expression and are the cis-regulatory sequences determining target genes’ spatiotemporal and quantitative expression. The reporter assays demonstrated that TR sequences enhanced KSHV inducible promoter activity more than x100 in 293T cells ^20^. Furthermore, TR strongly synergizes with the viral transcription activator, K-Rta, for the transactivation function ^20^; this makes the enhancer/promoter pair a very attractive gene element to develop KSHV-infection-specific gene therapy.

One of the critical barriers of gene therapy is the safe and efficient delivery of genetic material to the target tissues/cells, which is carried out by delivery vehicles. There are two gene therapy vectors: viral and non-viral ^21, 22^. Non-viral vectors comprise all the chemical and physical techniques and generally include chemical methods such as cationic liposome and synthetic polymers ^22, 23^, or physical methods such as gene gun ^24^, electroporation ^25^, ultrasound utilization ^26^, and magnetoreception ^27^. The advantages of non-viral vectors are their cost-effectiveness and less induction of immune reaction. However, the most successful gene therapy vectors available today are viral vectors, including retrovirus, adenovirus, lentivirus, and adeno-associated virus (AAV) ^28^. An advantage of viral vectors is higher delivery efficiency than non-viral methods. Viral genomes are modified for viral vectors by deleting some essential genes, which restricts their replication and allows safer gene delivery to the patients. Among the viruses used for gene therapy vectors, AAV was first discovered in 1965 as a co-infecting agent of adenovirus ^29^. The first infectious clone of AAV serotype 2 for human gene therapy was generated in 1982 ^30^. Since then, AAV serotypes 1-12 and over 100 AAV variants have been identified ^31^. AAV is a non-enveloped DNA virus with a genome of approximately 4.8 kbp single-stranded DNA ^32^. The coding capacity for the transgene is limited to ∼4.7 kbp, but this can be extended by splitting the transgene sequence into two viruses with co-infection ^33^. AAVs are naturally replication-deficient and require a helper virus for replication and dissemination ^34^. This self-replication deficiency and their ability to infect both dividing and non-dividing cells stably make them an excellent viral gene therapy vector. AAV gene therapy vectors have received FDA approval for commercialization - Luxturna for retinal dystrophy (NCT00999609) and Zolgensma for spinal muscular atrophy (NCT03306277). The number of FDA-approved AAV-associated drugs is expected to increase because many AAV gene therapy vectors are being tested at later clinical trial stages ^35^.

Indirect gene therapy, a method that converts the prodrug into a lethal (cytotoxic) drug within the tumor cells, demonstrates great promise and is being evaluated in clinical trials. The advantage of an indirect approach is to have an additional step, which makes the therapy adjustable and, therefore, safer for patients. The most studied indirect therapy is based on introducing the herpes simplex virus thymidine kinase (HSV-TK) gene with ganciclovir (GCV) administration ^36, 37^. The HSV-TK initiates the conversion of the antiviral drug, GCV, to a toxic metabolite, GCV-triphosphate, which inhibits DNA synthesis and induces cell apoptosis. The converted cytotoxic compound is also reported to have a bystander effect of killing surrounding non-transduced tumor cells, presumably due to released GCV-triphosphate from dying cells; this minimizes the necessity to transduce the vector to the 100% of tumor cells to be effective ^38, 39^.

Here, we designed and evaluated a KSHV-associated disease-specific gene therapy vector. We first examined the TK/GCV indirect therapy with AAV because KSHV is a herpesvirus, and GCV alone showed some efficacies in preventing KSHV-associated tumor progression in clinics ^40, 41, 42^. This report describes the construction and concept of the gene therapy vector and its initial characterization in the tissue culture model and the xenograft mouse model.

## Results

### Preparation and validation of KSHV-infection-specific gene expression cassette

Recent reports demonstrated that KSHV TR is not only a LANA binding sequence for KSHV episome maintenance but also possesses gene enhancer function ^18, 43^. Based on the findings, we devised an idea for a KSHV-tumor-specific gene therapy vector with an AAV. We designed the AAV vector encoding two copies of the TR sequence (**Figure 1a**), which should help maintain the transduced therapeutic vector in KSHV-infected (LANA expressing) cancer cells and increase KSHV inducible promoter activity. Because the TR enhances KSHV lytic gene promoter activity ^18, 43^, we cloned viral lytic gene promoter downstream of the TR sequence (**Figure 1b**). We selected the Ori RNA promoter (*OriP*), which is one of the K-Rta direct targets with higher promoter activity ^44, 45^, and the genomic fragments possess H3K27Ac and H3K4me3 active histone modifications in infected cells and localize proximity to TR in 3D genomic structure ^18, 44, 46, 47, 48^. We utilized an AAV transfer vector as a backbone to generate recombinant AAV for gene delivery. The procedure and vector design are depicted in **Figure 1a**. As described in more detail below, we cloned fluorescence protein (mCardinal) downstream of the Ori-RNA promoter (pAAV-TR2-*OriP*-mCardinal). The mCardinal was selected to distinguish the RFP signal produced from the r.219 KSHV viral genome ^49^ and monitor selective promoter activation in live cells.

**Figure 1.**
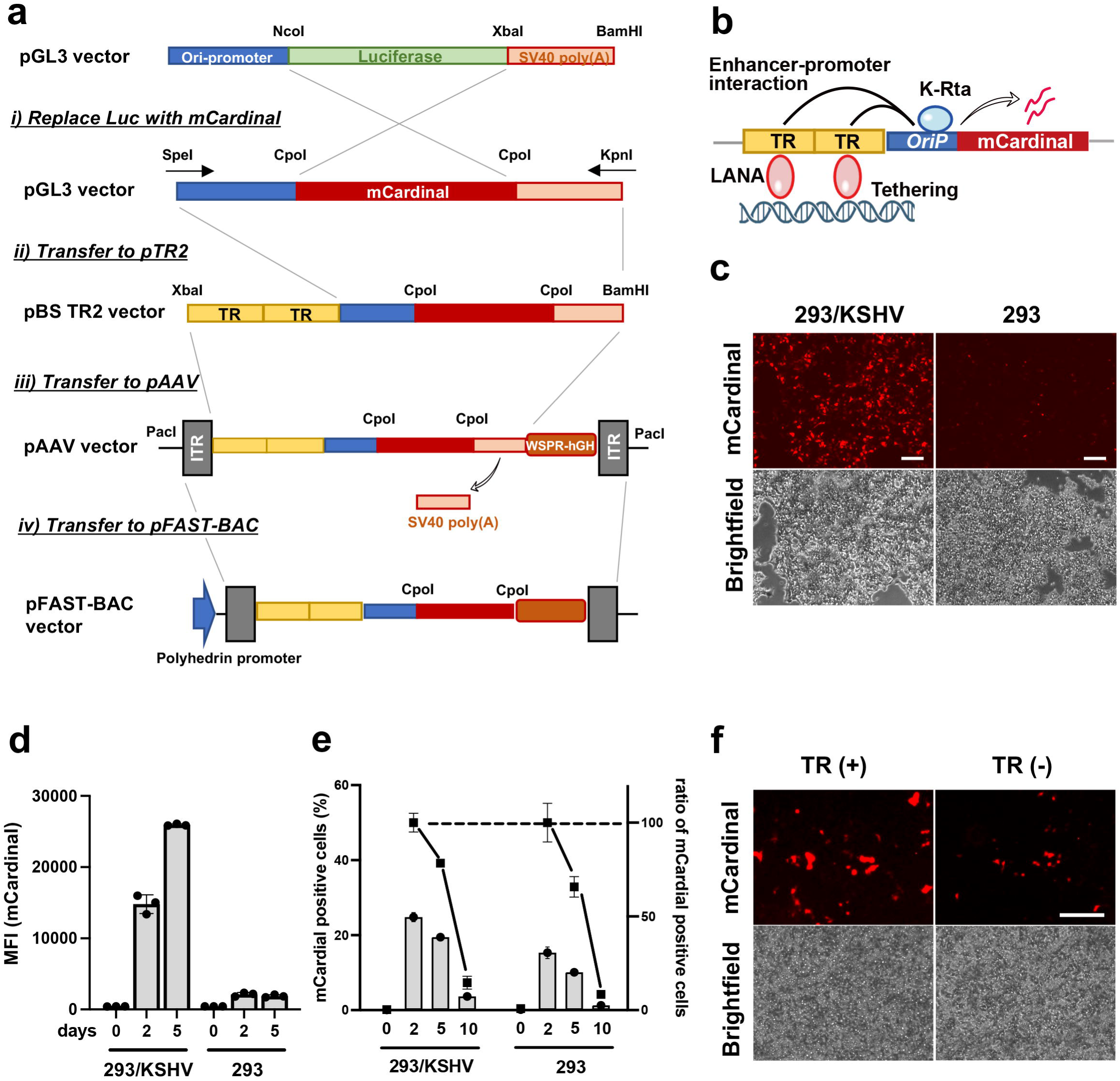
Construction of KSHV-infection-specific gene expression cassette. a. Schematic diagram of cloning strategy to generate specific gene therapeutic vector. Plasmid names are depicted in the left panel, and restriction enzyme sites used for cloning were also shown. TR: terminal repeat, ITR: inverted terminal repeat. b. Schematic representation of the regulatory mechanism of the gene cassette. The construct consists of terminal repeats (TR) and Ori RNA promoter (*OriP*) and the mCardinal fluorescent reporter gene. c. Fluorescent and bright field image of the 293/KSHV and the parental 293 cells. The pAAV-TR2-*OriP*-mCardinal vector was transfected into 293/KSHV and the parental 293 cells. Images were taken 48 hours post-transfection. Scales; 200μm.c. Mean fluorescence intensity (MFI). The mCardinal signal in 293/KSHV and the parental 293 cells after the transfection of the pAAV-TR2-*OriP*-mCardinal vector were measured with a Flow cytometer at 2 and 5 days post-transfection. e. The proportion of the mCardinal positive cells. The number of mCardinal expressing cells was determined by the Cy5 channel with the flow cytometry at indicated days after transfection. The relative proportion of mCardinal positive cells was depicted as the number of mCardinal cells at Day 2 as 100%. f. Enhanced exogenous gene expression with TR fragments. The pAAV-TR2-*OriP*-mCardinal vector with or without two copies of TR sequences was transfected to the 293/KSHV cells. Fluorescent and bright field images were taken two days after the transfection. Scales; 300μm

To examine if the assembled enhancer-promoter combination increases the exogenous gene expression in a KSHV infection-specific manner, we transfected the pAAV-TR2-*OriP*-mCardinal vector into KSHV r.219-infected 293 cells (293/KSHV cells) or parental 293 cells and monitored mCardinal expression. The signal intensity and the proportion of mCardinal positive cells were measured by flow cytometry. As expected, mCardinal intensity was approximately 15 times higher in 293/KSHV cells at 5 days post-transfection **(Figures 1c and d)**, and mCardinal signals were also maintained slightly longer in the 293/KSHV cells (**Figure 1e**). We also noticed that mCardinal signals were very weak in non-KSHV infected cells, suggesting that KSHV-infection, presumably, K-Rta protein expression from infected KSHV genomes, enhanced mCardinal expression. The effects of the TR sequence on enhancing gene expression were further confirmed by transfecting pAAV-*OriP*-mCardinal vector with or without TR2. Consistent with the previous studies ^20^, the vector with TR showed brighter signals in the 293/KSHV cells **(Figure 1f)**.

### Preparation of recombinant AAV

To conveniently and cost-effectively prepare a large scale of recombinant AAV in-house, we next adapted the baculovirus-based AAV production platform. With serum-free defined culture media, we could also avoid using animal proteins, which reduces biosafety concerns. AAV therapeutics prepared with recombinant baculovirus platforms are being evaluated in human clinical trials ^50^.

The TR2-*OriP*-mCardinal fragment was first moved into the baculovirus transfer vector as described in material methods (**Figure 1a**), and the recombinant baculovirus, which carries the entire AAV transfer genome, including both inverted repeat sequences, was generated. The insect cells were then co-infected with the recombinant baculovirus expressing Rep78, Rep52, and AAV8 capsid proteins. Baculovirus, a large DNA virus, serves as a helper virus for AAV in insect cells, producing AAV virion with the transfer DNA. Co-infected Sf9 cells were harvested 72 hours post-infection. Recombinant AAV8 (AAV8-TR2-*OriP*-mCardinal) was purified with iodixanol gradient ultracentrifugation for isolation (**Figure 2a**). With the baculovirus-mediated AAV preparation, we routinely isolated approximately 2 mL of 10^13^ copies/ml of recombinant AAVs from 100 mL of Sf9 suspension culture. The purity of AAVs was monitored by SDS-PAGE gels with capsid protein bands as indicators, and highly pure fractions (fractions 1-4) were combined for use in experiments (**Figure 2b, left**). The frequencies of empty capsids that would impair transduction efficiency were also monitored by electron microscopy (**Figure 2c**). The result showed that more than 90% of purified AAV virions contained transfer DNAs.

**Figure 2.**
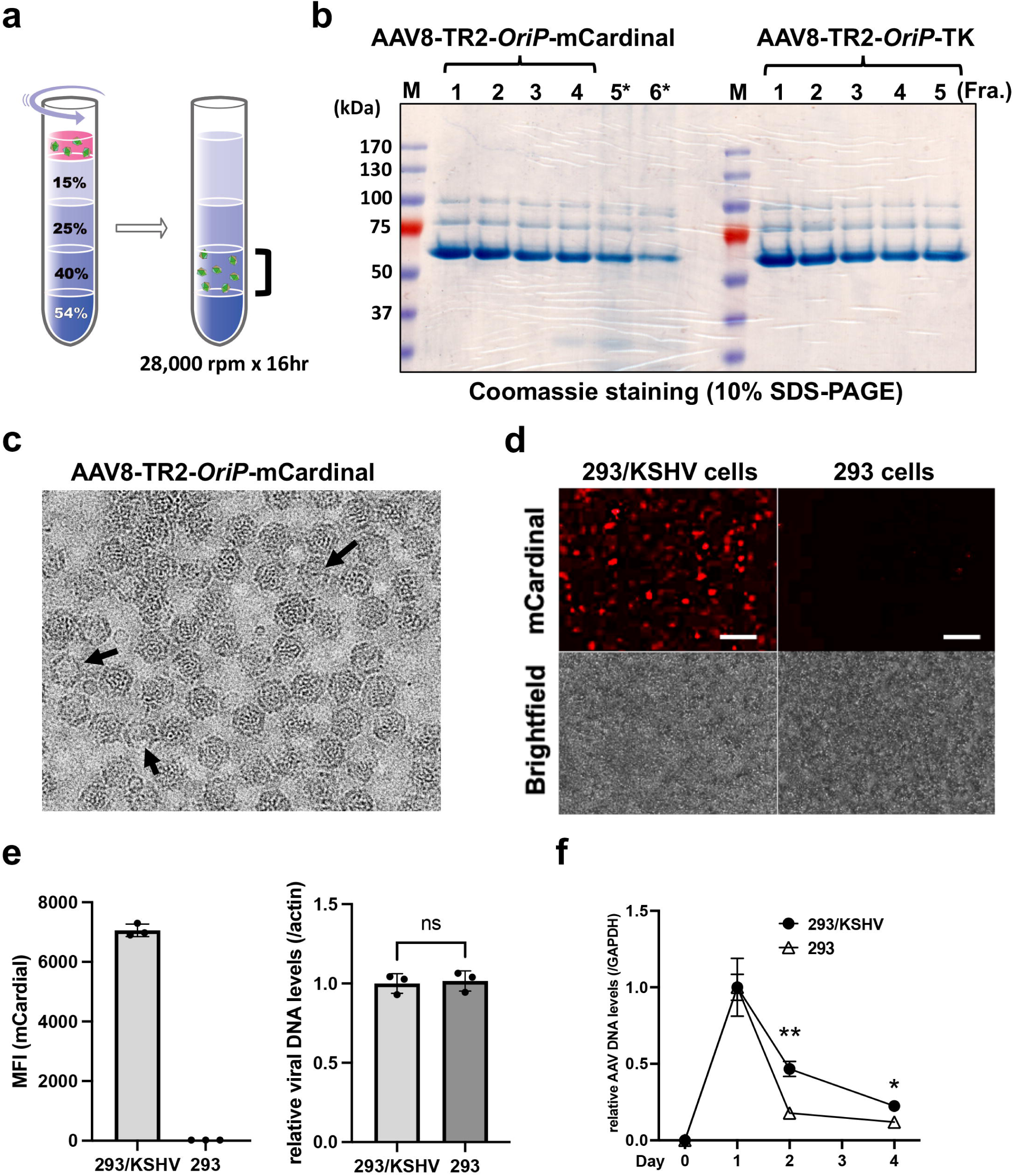
Activation of TR2-*OriP* promoter in KSHV infected cells. **a. Schematic diagram of the iodixanol gradient ultracentrifugation for AAVs isolation.** The fraction with 40% iodixanol contains AAVs. b. SDS-PAGE gels. The 40% iodixanol layer was fractionated from bottom to top 1 to 6. Coomassie staining shows the AAV capsid protein, VP1, VP2, and VP3. The fractions 1 to 4 for AAV8-TR2 *OriP* mCardinal were pooled, concentrated, measured DNA copies, and used for the following studies. The molecular size marker is indicated on the left side of the gel. **c. Transmission electron microscopy (TEM)** The representative TEM images for the AAV8-TR2-*OriP*-mCardinal is shown. Putative empty capsids are marked with an arrow. **d Fluorescent and bright-field images.** The 293/KSHV and the parental 293 cells were infected with AAV8-TR2-*OriP*-mCardinal and images were taken 72 hours post-transduction. Scales; 100 μm **e. Mean fluorescence intensity (MFI) of the mCardinal signal and AAV genome copies.** MFI was measured with a flow cytometer, and AAV DNA copies in transduced cells were determined by qPCR. Relative AAV DNA levels were measured at the indicated days after infection. GAPDH coding sequence was used for internal control. Data was analyzed using a two-sided unpaired Student’s t-test and shown as mean ± SD. **f. Relative abundance of AAV DNA copies.** AAV DNA copies were measured by qPCR and compared between the 293/KSHV and the parental 293 cells. Twenty-four hours after AAV-mCardinal transduction in 293/KSHV cells was designated as 1. GAPDH coding sequence was used for internal control. Data was analyzed using a two-sided unpaired Student’s t-test and shown as mean ± SD.

With purified recombinant AAVs in our hand, we next transduced the AAV8 TR2-*OriP*-mCardinal to 293/KSHV cells or the parental 293 cells and examined the amount of mCardinal expression with brightness. In AAV-transduced cells, the single-stranded AAV transfer genomes become double-stranded circular DNA, like KSHV episome. Consistent with plasmid transfection (**Figures 1c and d**), mCardinal signals were significantly brighter in 293/KSHV cells than parental 293 cells (**Figure 2d and e, left**). To rule out the possibility that the AAV-infection efficiency is different between KSHV-infected and non-infected cells, we measured intracellular AAV DNA after infection. The results confirmed no differences in AAV infectivity between KSHV-infected and parental 293 cells **(Figure 2e, right)**. Flow cytometry analyses also showed that transduced AAV genomes were maintained better in 293/KSHV cells (**Figure 2f**). These results suggested that AAV8-TR2*-OriP* can selectively express exogenous genes in a KSHV-infection-specific manner.

### Selection of thymidine kinase gene

Having vectors that preferentially express the exogenous genes in KSHV-infected cells, we next replaced mCardinal with a therapeutic gene (**Supplementary** Figure 1a). We selected the thymidine kinase (TK)/ ganciclovir (GCV) for an indirect gene therapy. This is because KSHV is a herpesvirus, and GCV alone has been shown to control KSHV-associated tumor progression ^40, 41, 42^. Having a step for the conversion of the prodrug into a cytotoxic drug within the tumor cells makes the therapy adjustable. The converted cytotoxic compound is also reported to have a bystander effect of killing surrounding non-transduced tumor cells (**Supplementary** Figure 1b); this minimizes the necessity to transduce the AAV8-TR2-*OriP*-TK to 100% of tumor cells to be effective ^38, 39^. We employed TKSR39, an engineered form of TK, which improved GCV conversion approximately ten times ^51^. The codon optimization for humans has also been reported to increase cell killing ^52^. Accordingly, we designed and synthesized the modified TK gene and purified recombinant AAVs with the baculovirus-based AAV production platform (Figure 2b, right).

### AAV8-TR2*-OriP*-TK with Ganciclovir (GCV) induced cancer cell death in a KSHV infection-specific manner

Next, we examined the degree to which the transduction of the AAV8-TR2*-OriP*-TK specifically induces KSHV-infected cell death and spares non-KSHV-infected cells. Preserving non-KSHV-infected cells should minimize side effects and increase the therapeutic window. We first used two KSHV-infected cell lines (293 cells and iSLK cells) and parental non-infected cells to consider the cell line bias. KSHV-infected (+) and non-infected (-) cells were transduced with AAV8-TR2-*OriP*-TK. Twenty-four hours post-transduction, we incubated with GCV (5 μM) with or without OTX015 (BRD4 inhibitor). Previous reports showed that the BRD4 inhibitor induces KSHV reactivation^53, 54, 55, 56^. Therefore, we expected OTX015 would increase TK expression from AAV therapeutic vectors. The results showed that transduction of AAV8-TR2*-OriP*-TK induced cell death only in the presence of GCV in both 293 and iSLK cells (**Figures 3a and c**). More importantly, the presence of latently infected KSHV strongly sensitized to the AAV8-TR2-*OriP*-TK (**Figures 3b and d**). In the case of 293 cells, the presence of KSHV is necessary to induce cell death with the AAV8-TR2*-OriP*-TK and GCV; the results suggest that the TR2-*OriP* activation is strictly regulated by KSHV infection (**Figures 3a and b**). Decreased cell viability in iSLK cells without KSHV infection may be due to leaky expression of K-Rta from exogenous K-Rta cassette, and may weakly activate TK expression from the AAV vector. **Figure 3e** shows the morphology of the iSLK/KSHV and iSLK cells 2 days after AAV8-TR2*-OriP*-TK infection. The histone deacetylase inhibitor, suberoylanilide hydroxamic acid (SAHA), is known to trigger KSHV reactivation ^37, 56, 57, 58^, similarly as OTX015. We expected the anti-cancer drugs that stimulate KSHV reactivation and, therefore, stimulate TR2-*OriP* activity to enhance cancer cell killing. OTX015 or SAHA were then incubated with GCV to examine the synergistic cell killing and proliferation. As we expected, SAHA and OTX015 enhanced TR2-*OriP* activity (**Supplementary** Figure 2a) and inhibited KSHV-infected cell growth (**Supplementary** Figure 2b **and Supplementary Movies 1a-d**).

**Figure 3.**
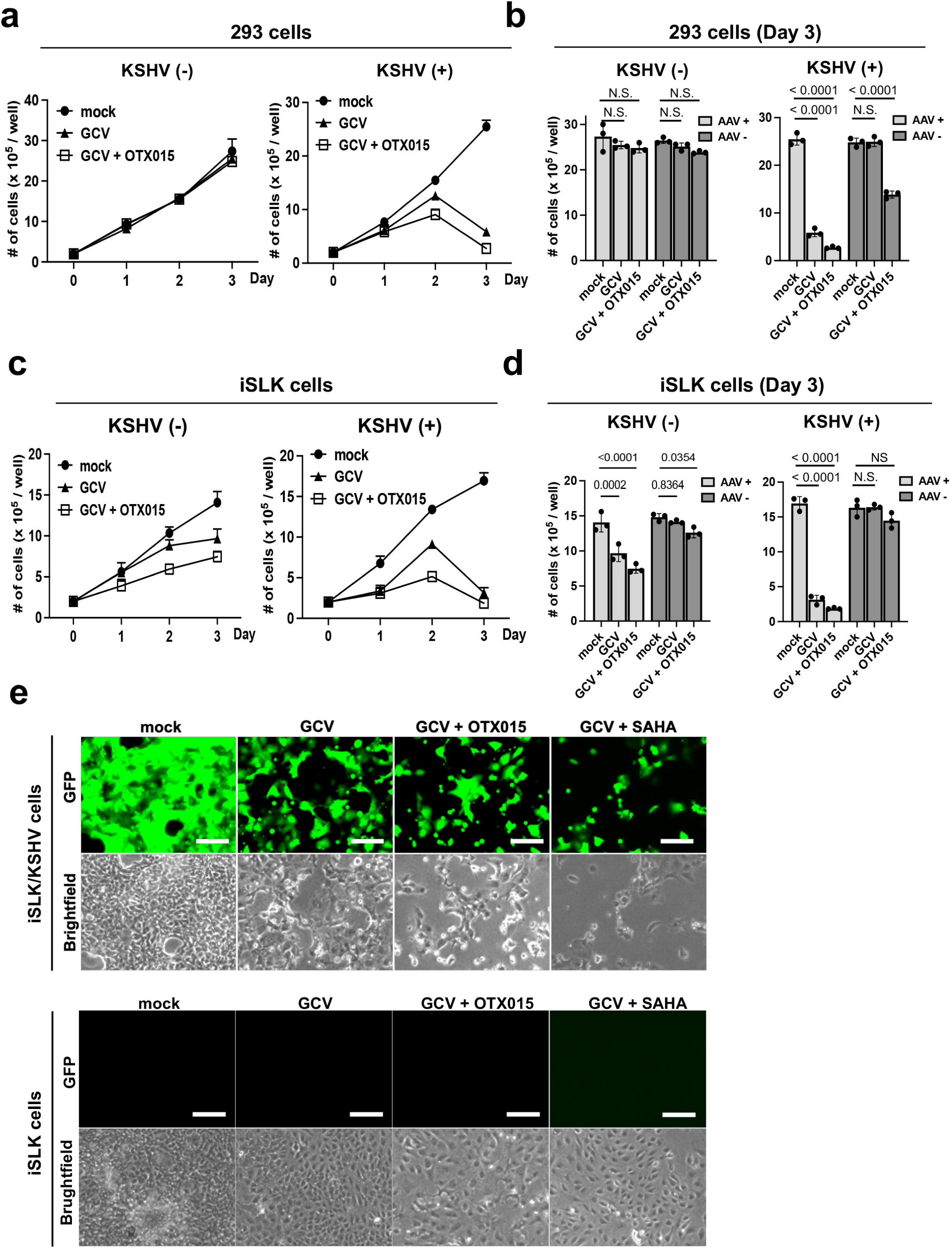
Inhibition of cell growth by AAV8-TR2-*OriP*-TK in KSHV infection-specific manner. **a. 293 cells growth.** 293 cells or KSHV-infected 293 cells were seeded in 12 well plates and transduced with AAV8-TR2-*OriP*-TK. Cells were treated with mock, GCV (5 μM), or GCV and OTX015 (200 nM). Live 293 cells were counted every day for three days. The total number of cells in each well was counted in triplicate, and a bar graph was generated with mean ± SD. **b. Bar chart with or without AAV8-TR2-*OriP*-TK transduction in 293 cells.** Live 293 and 293/KSHV cells at Day 3 with or without AAV8-TR2-*OriP*-TK are shown in bar charts. Data was analyzed using a two-sided unpaired Student’s t-test and shown as mean ± SD. **c. iSLK cells growth**. iSLK cells or KSHV-infected iSLK cells were seeded in 12 well plates and transduced with AAV8-TR2-*OriP*-TK. Cells were treated with mock, GCV (5 μM), or GCV and OTX015 (200 nM). The total number of cells in each well was counted in triplicate, and a bar graph was generated with mean ± SD. **d. Bar chart with or without AAV8-TR2-*OriP*-TK transduction in iSLK cells.** Live iSLK and iSLK/KSHV cells at Day 3 with or without AAV8-TR2-*OriP*-TK are shown in bar charts. Data was analyzed using a two-sided unpaired Student’s t-test and shown as mean ± SD. **e. Fluorescent and bright field images.** iSLK cells with or without KSHV infection were transduced with AAV8-TR2-*OriP*-TK. Cells were treated with the indicated drug combination. Images were taken three days after treatment of OTX015 (200 nM) or SAHA (1 μm). Scales; 200μm

### AAV8-TR2-*OriP*-TK specifically inhibits the growth of KSHV-infected ECFCs

While 293 and iSLK cells are important research tools for KSHV infection and report differences made by KSHV infection to the sensitivity to the vector, those kidney cells are unlikely to be natural target cells in *vivo*. A recent study suggested that endothelial colony-forming cells (ECFCs) may be the origin of KS ^59, 60, 61^. To examine if our vector is also effective in KSHV-infected ECFCs, we differentiated induced pluripotent stem cells (iPSCs) into ECFCs by following previous studies ^62^ **(Figure 4a)**. The differentiation of ECFCs was confirmed by the upregulation of ECFC markers, such as CD34, Prox-1, Flt-4, and LYVE-1, while the expression of the pluripotency markers, Oct3/4, Nanog, and Sox2, were significantly reduced when we triggered cell differentiation **(Figure 4b)**. Consistent with previous studies, KSHV efficiently infected ECFCs but not iPSCs ^59^ **(Figure 4c)**. KSHV-infected ECFCs or parental ECFCs were transduced with AAV8-TR2-*OriP*-TK and treated with GCV (5 μM) the following day. Live cells were counted for three days. The results showed that AAV8-TR2-*OriP*-TK with GCV strongly induced KSHV-infected cell death, whereas uninfected ECFCs were not affected **(Figure 4d)**. Fluorescent and bright field imaging further confirmed that induction of cell death strictly depended upon KSHV infection and GCV **(Figure 4d, e)**.

**Figure 4.**
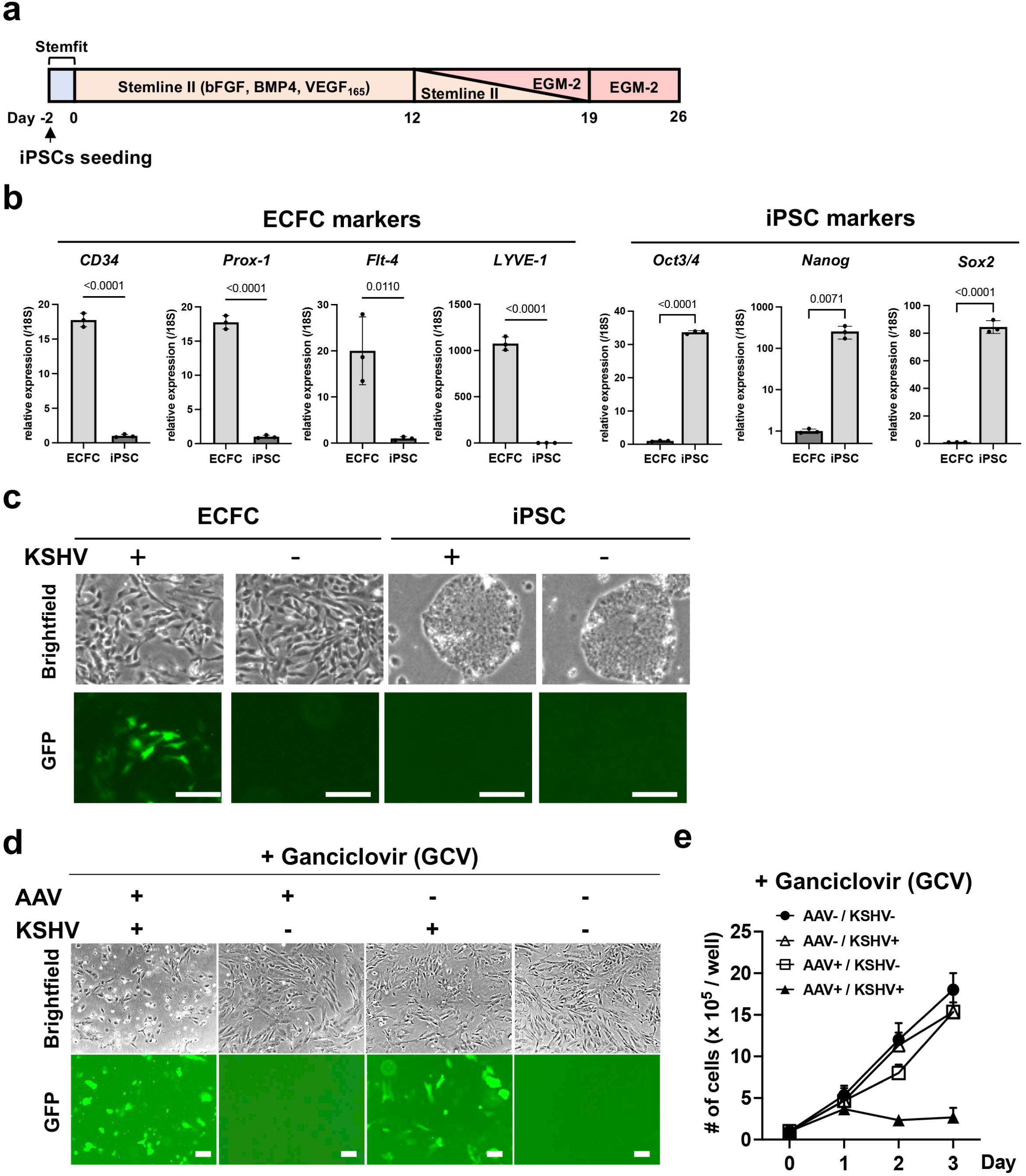
AAV8-TR2-*OriP*-TK selectively Inhibits KSHV-infected ECFCs growth. **a. Schematic diagram of ECFCs differentiation from iPSCs.** iPSC differentiation was induced by bFGF, BMP4, and VEGF_165_ in the Stemline II media, followed by EGF-II culture. **b. Relative gene expression of iPSCs and ECFCs (Day 26).** CD34, Prox-1, Flt-4, and LYVE-1 genes were used as differentiation markers for ECFCs, while the Oct3/4, Nanog, and Sox2 genes were used as reprogramming markers. 18S was used for internal control. Data was analyzed using a two-sided unpaired Student’s t-test and shown as mean ± SD. **c. Fluorescent and bright field cell images.** r.219 KSHV (MOI = 1) was infected to differentiating cells at 12 days post induction of iPSC differentiation or parental iPSCs. Images were taken 14 days after infection. **d. Fluorescent and bright field cell images.** ECFCs or KSHV-infected ECFCs transduced with AAV8-TR2-*OriP*-TK were treated with GCV (5 μM), and images were taken three days after GCV treatment. **e. Cell growth.** ECFCs or KSHV-infected ECFCs were seeded in 6 well plates and transduced with AAV8-TR2-*OriP*-TK. The following day, cells were treated with DMSO or GCV (5 μM). Live ECFCs were counted every day for three days. The total number of cells in each well was counted in triplicate, and a bar graph was generated with mean ± SD.

### AAV8-TR2-*OriP*-TK prevents KSHV replication during reactivation

One of the benefits of using GCV is that it inhibits KSHV DNA replication in reactivating cells, therefore preventing reactivated KSHV from infecting neighboring cells. To examine if AAV8-TR2*-OriP*-TK synergizes with endogenous KSHV TK and inhibits KSHV replication in the presence of GCV, we reactivated KSHV from iSLK cells with a combination of doxycycline with sodium butyrate and measured the effects of transduced AAV8-TR2-*OriP*-TK on KSHV replication (**Figure 5a**). This tactic is similar to delivering a “*Trojan’s horse*” to the KSHV residence. The results showed that in the delivery of AAV8- TR2-*OriP*-TK in KSHV-infected cells, GCV abolished K8.1 late gene expression more than 100-fold, while PAN-RNA expression decreased approximately 10-fold (**Figure 5b**), which is consistent with the weaker RFP signal in AAV8-TR2-*OriP*-TK infected iSLK/KSHV cells (**Figure 5a**). The late gene K8.1 expression depends on the viral DNA replication, suggesting that KSHV DNA replication was significantly diminished. KSHV virion production in culture media was also reduced approximately 10-fold with additional AAV8-TR2-*OriP*-TK (**Figure 5c**). Consequently, AAV8-TR2-*OriP*-TK prevents KSHV infections from 90% to 7% when we transfer the culture supernatant to freshly prepared iSLK cells (**Figure 5d, e**). These results suggested that AAV8-TR2*-OriP*-TK selectively inhibited KSHV-infected cell growth and prevented the spreading of KSHV infection from reactivation.

**Figure 5.**
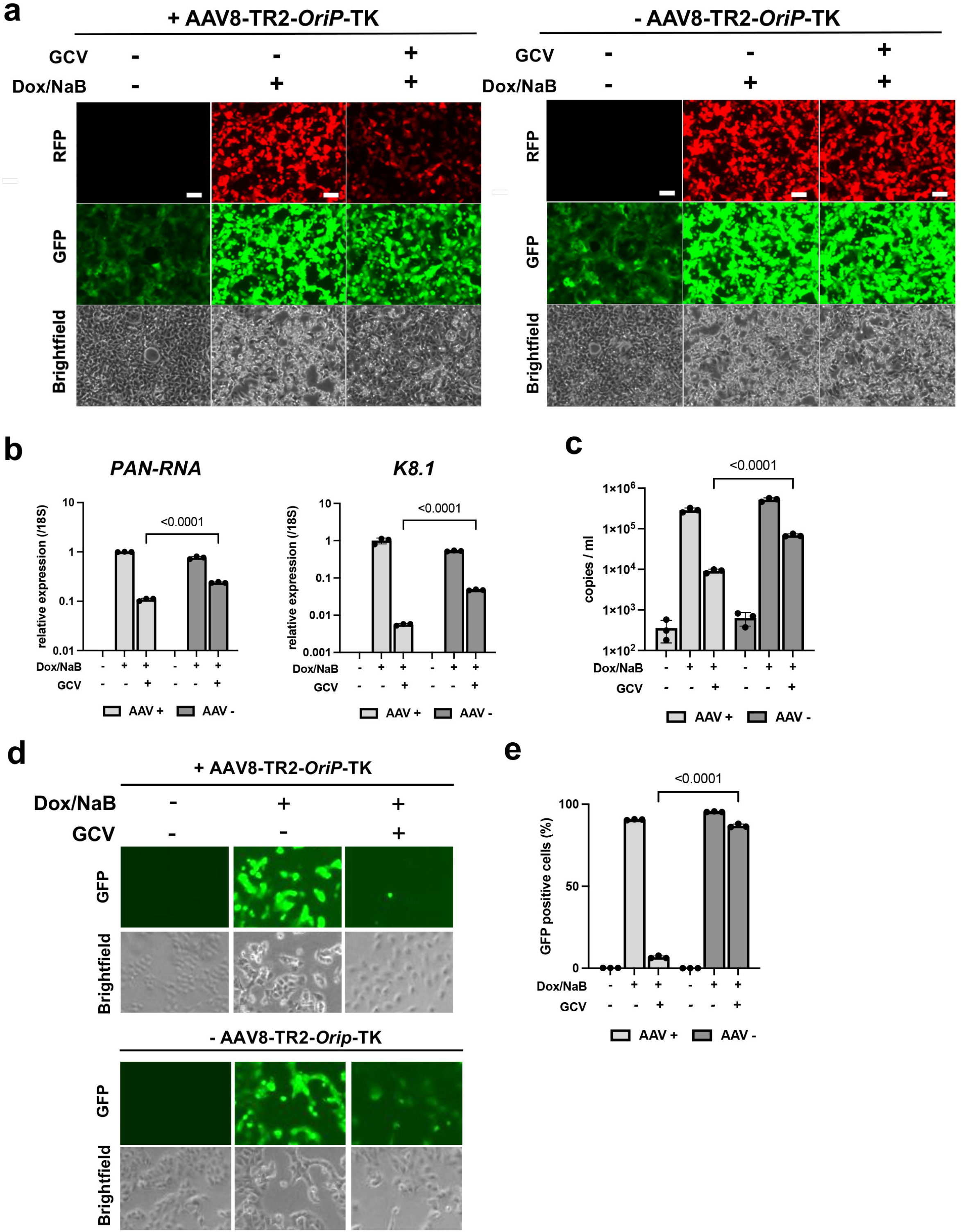
AAV8-TR2-*OriP*-TK prevents KSHV reactivation and re-infections. **a. Fluorescence and bright field cell images.** iSLK/KSHV cells were treated as indicated, and the cells were imaged with fluorescence microscopy at 48 hours post-stimulation. AAV8-TR2-*OriP*-TK transduction with GCV treatment decreased the RFP signal, an indication of decreased PAN-RNA promoter activation. **b. Viral gene expression.** PAN-RNA or K8.1 transcripts were measured by RT-qPCR with specific primer pairs. Transcripts were normalized with 18S rRNA. The relative transcripts in AAV8-TR2-*OriP*-TK transduced without GCV were normalized as 1. Data was analyzed using a two-sided unpaired Student’s t-test and shown as mean ± SD. **c. Capsidated viral DNA copies.** The KSHV virion copy number with or without reactivation in the presence or absence of GCV (5 μM) was measured with qPCR. KSHV virions were collected from the culture supernatant 4 days post-reactivation. Data was analyzed using a two-sided unpaired Student’s t-test and shown as mean ± SD. **d. KSHV infection to freshly prepared iSLK cells.** The KSHV infection with reactivated virus was evaluated by infecting freshly prepared iSLK cells**. 1 ml** culture supernatant 4 days post-reactivation in the presence or absence of GCV (5 μM) was mixed with freshly prepared iSLK cells. Images were taken two days after infection. **e. Flow cytometry.** Frequencies of KSHV infection were measured with flow cytometry. Data was analyzed using a two-sided unpaired Student’s t-test and shown as mean ± SD.

### Bystander effect of AAV8-TR2-*OriP*-TK in KSHV-infected cell population

One of the bottlenecks of gene therapy, in general, is the efficacy of gene delivery. Even with the topical transduction of AAV8-TR2-*OriP* TK to KS skin or oral KS lesions, we may not achieve 100% transduction efficacies. Thus, having the bystander cell-killing effect is critical. To monitor the bystander cell killing, we first prepared a recombinant BAC16 virus, which constitutively expresses the mCherry-H2B gene under the EF1alpha promoter. We replaced EGFP genes encoded in the BAC backbone with the mCherry-H2B coding sequence. The iSLK/KSHV BAC16 cells (EGFP-positive) were transduced AAV8-TR2-*OriP*-TK and co-cultured with iSLK/mCherry-KSHV cells (without AAV-transduction) at 1:1 ratio, followed by GCV treatment for three days **(Figure 6a)**. The results showed that GCV incubation induced cell death not only in EGFP-positive cells (AAV-infected) but also in mCherry-positive cells (bystander cells), demonstrating the bystander effect of AAV-TK infection (**Figure 6b**).

**Figure 6.**
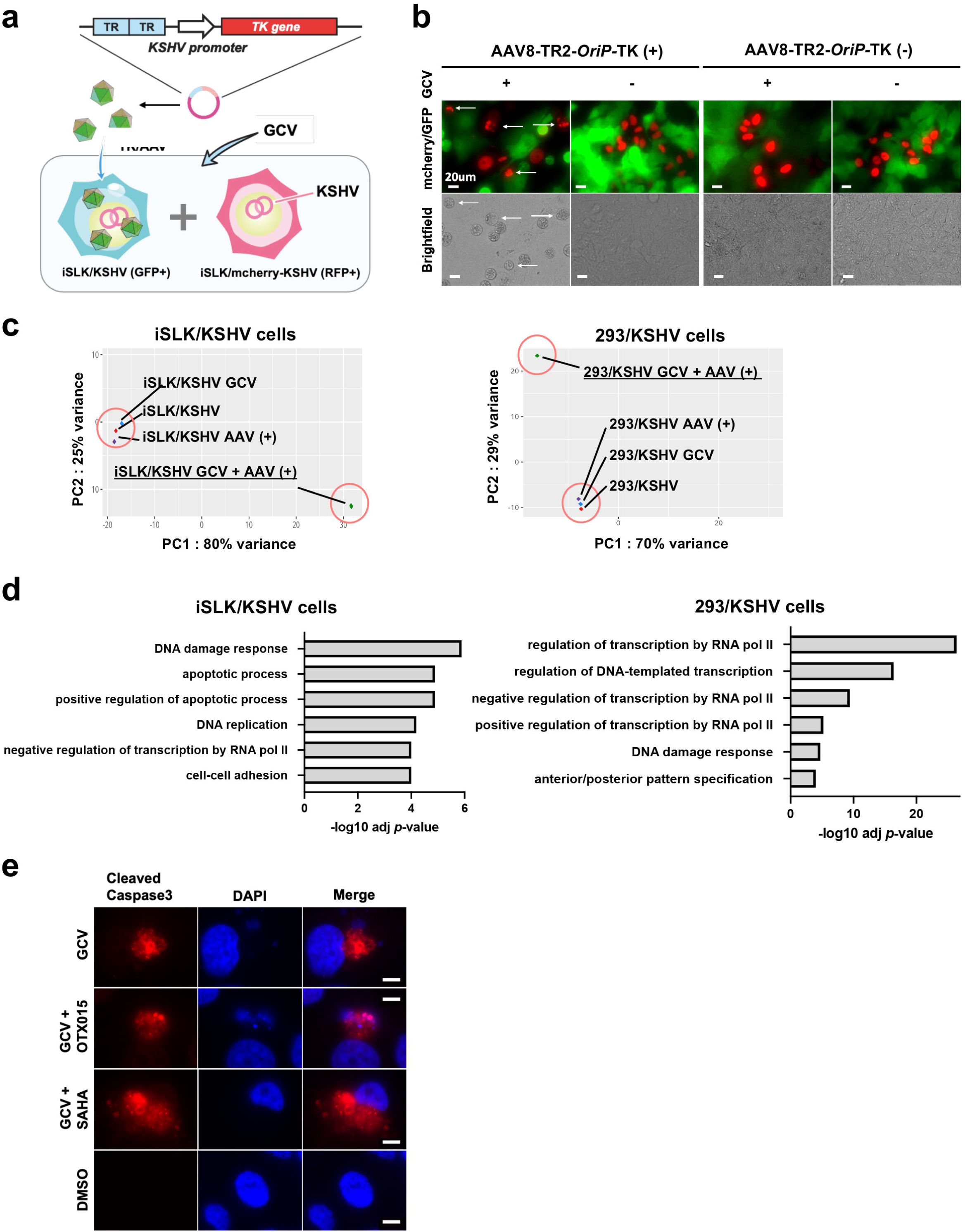
Bystander effects and transcription profiles. **a. Schematic diagram of study design for the bystander effect.** Two types of KSHV-infected iSLK cells were prepared (Green and Red), and AAV8-TR2-*OriP*-TK was transduced only to EGFP-positive cells. Green and red (mCherry-positive) cells were mixed with a 1:1 ratio (10^5^ cells each) and cultured for four days in the presence of GCV (5 μM). Bystander effects were assessed with the viability of red cells. **b. Fluorescence cell images.** Arrows indicate dead red cells determined by cell morphology and fractured nucleus. Images were taken four days after the mixture of iSLK/KSHV (green) and iSLK/mcherry-KSHV (red) cells **c. Principle component analyses.** Total RNA sequence was performed with iSLK/KSHV and 293/KSHV cells. Cells were treated as indicated in the panel. **d. Pathway analyses.** Cellular signaling pathways that are significantly altered in GCV with AAV8-TR2-*OriP*-TK transduction were shown. The results indicated a strong induction of DNA damage responses with subsequent induction of cell apoptosis. **e. Immunofluorescence assays.** AAV8-TR2-*OriP*-TK transduced iSLK/KSHV cells were stained with antibody specific to cleaved caspase 3 and rabbit-647 secondary antibody. Cleaved caspase 3 expression in iSLK/KSHV cells under GCV (5 μM) treatment was confirmed with immune staining.

### AAV8-TR2-*OriP*-TK with GCV induces cell apoptosis

We performed transcription profiling to enhance cancer cell killing further when needed. The total RNA was extracted after 48 hours of GCV incubation, and RNA-sequence was performed. PCA analysis demonstrated that AAV8-TR2-*OriP*-TK infection showed very similar transcriptional profile in the absence of GCV in iSLK/KSHV and 293/KSHV cells (**Figure 6c)**. Two pairs of cell lines were used to identify the signaling pathways commonly regulated by AAV8-TR2-*OriP*-TK under GCV. DAVID analysis for comprehensive functional annotation ^63^ showed that DNA damage response and apoptotic process were enriched in iSLK cells transduced with AAV8-TR2-*OriP*-TK with GCV, while DNA damage response was the most enriched in 293 cells (**Figure 6d)**. We confirmed the presence of cleaved caspase 3 with GCV in AAV8-TR2-*OriP*-TK transduced cells **(Figure 6e)**. These results suggest that AAV8-TR2-*OriP*-TK induced cell apoptosis with GCV via induction of DNA damage. Consistent with the result, induction of cell apoptosis with GCV was also reported in other cell types ^64, 65^.

### Anti-tumor activity in xenograft mice

We finally examined the efficacy of cancer cell killing over toxicities in a xenograft mouse model. We first infected KSHV-infected iSLK cells with AAV8-TR2-*OriP*-TK in the culture dish and expanded the cells. Four days after AAV8-TR2-*OriP*-TK transduction, iSLK cells were harvested, and 5 x10^6^ cells were implanted at the subcutaneous of male NRG mice (right hind leg). No-AAV transduced cells were used as a comparison (left hind leg). GCV or PBS (50 mg/kg, twice a day) was administrated intraperitoneal (IP) for five days starting from two days after iSLK cells SQ injection (**Figure 7a**). As shown in **Figure 7b**, tumor mass was significantly diminished with GCV treatment. The tumor growth inhibition was AAV8-TR2-*OriP*-TK transduction and GCV administration specific because iSLK/KSHV cells without AAV8-TR2-*OriP*-TK planted in the same mice with GCV continued to grow. Similarly, without GCV treatment, AAV8-TR2-*OriP*-TK failed to inhibit tumor growth (**Figures 7c, d, e**). As expected with a clinical drug, the GCV-treated mice did not show signs of discomfort or weight loss during the experiment periods. Immunohistochemistry with Ki67 antibody showed that AAV8-TR2-*OriP*-TK with GCV strongly inhibited iSLK cell growth, and the overall number of KSHV-infected live cells (EGFP-positive) in the tumor mass is also lower **(Figure 7f)**. Together, these results demonstrated the efficacies of AAV8-TR2-*OriP*-TK as a monotherapy and as an adjuvant to enhance the efficacies of chemotherapy drugs that induce KSHV reactivation.

**Figure 7.**
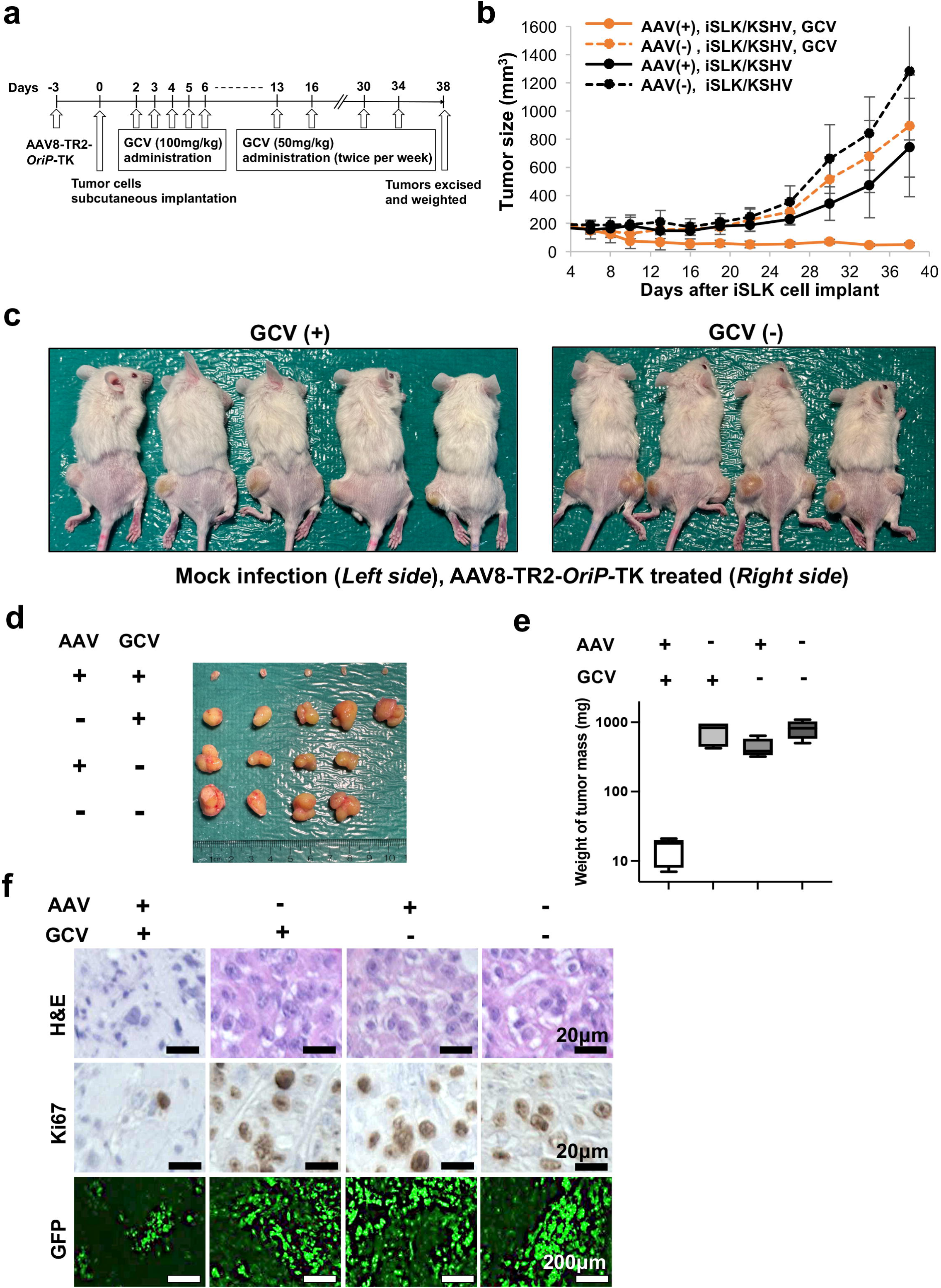
AAV8-TR2-*OriP*-TK inhibits tumor growth in a xenograft model. **a. Schematic diagram of treatment schedules.** KSHV-infected iSLK cells were transduced with AAV8-TR2-*OriP*-TK and xenograft subcutaneous. AAV8-TR2-*OriP*-TK transduced iSLK cells were implanted into the right hind leg of the mice, while non-transduced iSLK cells were injected into the left hind leg. GCV was administered as indicated. **b. Measurement of tumor sizes.** Tumor volumes (mm^3^) were measured every 4 days. **c. Mouse images at Day 38. d. Images of tumor mass extracted from subcutaneous. e. Tumor volumes.** The weight of tumor mass (mg) were plotted, and treatments were indicated at the top of the panel. **f. Hematoxylin-eosin (H&E) staining and Immunohistochemistry (IHC) staining of Ki-67 in mouse tissue sections.** Representative images of IHC stainings and EGFP signals found in individual tumors. Scales: 20μm (top and middle), 200μm (bottom).

## Discussion

The incidence of KS in HIV+ patients has decreased since the introduction of antiretroviral therapy; however, KS remains the most common HIV-associated malignancy ^66^. Cancers with viral etiology, like KS, have more apparent therapeutic targets because the malignant cells usually express viral proteins, and those viral proteins are also responsible for cancer cell growth. Rapid LANA protein knock-down also led to the elimination of the KSHV genome from the cancer cells ^67^. However, finding small molecules to target LANA function effectively is still a work in progress in the KSHV research community ^10, 11^.

Instead of targeting LANA directly, we took advantage of LANA function and well-developed KSHV transcription program to target KSHV-infected cells ^68, 69^. The idea is based on a decade of KSHV gene regulation studies that showed that KSHV appears to establish a transcription program, which minimizes effects from cellular signaling events via well-designed enhancer and promoter DNA sequences that are primarily regulated by KSHV proteins ^20, 44, 46, 69^. The relatively “closed” transcription regulatory mechanism designed for KSHV proteins allowed us to generate a gene expression vector whose expression is restricted in KSHV-infected cancer cells. Based on previous KSHV promoter screening, we selected the Ori-RNA promoter, which is a direct target of K-Rta ^20, 44, 46, 70^. We also showed that the Ori-RNA promoter is recruited by the LANA protein complex, presumably regulated by the LANA protein complex assembled at the TR region (enhancer region) ^18^. Accordingly, by physically neighboring two genomic regions (TR and Ori-RNA promoter) by design, we expected to argue/tighten the LANA/TR-mediated transcription regulation, which we expected to make the transcription regulation more specific to KSHV infection. We were initially afraid that the Ori-RNA promoter, although the strongest among KSHV lytic inducible promoters at the basal level, is still not strong enough to drive therapeutic gene expression to the level that induces cancer cell death. However, the weaker promoter worked positively by sparing non-KSHV-infected cells. Nonetheless, having several building blocks that include CMV gene enhancer element, other KSHV lytic gene promoters, and selection of therapeutic genes, we should be able to generate multiple therapeutic vectors flexibly.

To evaluate our gene therapy approach, we first searched for cell lines that could infect both AAV8 and KSHV effectively and that were also used widely in the KSHV research community. The screening found that SLK cells and 293 cells were susceptible to KSHV and AAV8 infections. Later, we confirmed the efficacies of the approach in more clinically relevant ECFC ^59^. However, we also found that the KSHV naturally-infected primary effusion lymphoma cells (BC1, BC3, and BCBL-1) were refractory to AAV8 infection, suggesting that additional capsid engineering will be required when we target PEL cells ^71, 72, 73^. Accordingly, our vector is primarily for skin and oral KS. Targeting skin and oral KS also has another important advantage for gene therapy; delivery. We should be able to administrate topically with micro-needles or band-aids with hydro-gels. Alternatively, using mini-plasmid DNAs may also reduce cost significantly and increase the shelf life of the drug.

We demonstrated two important advantages of applying the AAV8-TR2*-OriP*-TK to the KSHV-associated malignancies: cancer cell killing and inhibition of KSHV replication from reactivating cells. Importantly, GCV alone has already been proven to control KSHV-associated disease progression in clinics, presumably due to the inhibition of spreading KSHV from spontaneous reactivation, which also prevents inflammatory cytokine expressions ^74, 75, 76^. We expect that the additional TK expression from AAV8-TR2*-OriP*-TK in KSHV-infected cancer cells should enhance the GCV effects locally and further widen the therapeutic windows to control KSHV-associated diseases.

Several anti-cancer drugs are known to reactivate KSHV in latently-infected cancer cells ^53, 56, 77^. The oncolytic strategy, which stimulates KSHV reactivation via an anti-cancer drug to synergistically kill cancer cells, has been examined in clinical trials^77^. The hurdle of the oncolytic approach is to reactivate KSHV in all of the latently infected cancer cells. Transient expression of K-Rta with the stimuli does not always lead to a complete cycle of KSHV lytic replication ^19, 78^. We showed the enhancement of cancer cell killing with the AAV8-TR2-*OriP*-TK and bystander effects with transduction of the vector in 50% of the cell population in the dish. We expect that having the vector as an adjuvant should improve the outcome of the chemotherapy, especially for oncolytic therapy, which relies on small molecular drugs to stimulate K-Rta expression in the cancer cells. Further studies will identify the most effective combinations with AAV-TR2-*OriP*-TK.

The xenograft studies evaluated the efficacies of tumor cell killing versus toxicity of the gene therapy vector in mouse patients. We transduced AAV to cancer cells prior to engrafting in mice, which minimized the toxicity of systemic AAV transduction to normal mouse tissues. Because we found no decreased cell viability in non-KSHV infected 293 cells or ECFCs with AAV8-TR2-*OriP*-TK with GCV, we expect little toxicity by the AAV8-TR2*-OriP*-TK systemic infection. However, monitoring liver damage after the systemic injection (IV) should still be needed because AAV8 is known to infect a majority of tissues, especially in the liver ^79^. We hope to complete pre-clinical studies, including comprehensive toxicology and biodistribution analyses of AAV8-TR2*-OriP*-TK, and file an Investigator New Drug. By following the paths for AAV-based drugs that have already been approved by the FDA or are in the late stages of clinical trials, we hope to move on to clinical trials relatively smoothly.

In summary, based on a decade of basic KSHV gene transcription studies, we designed, developed, and demonstrated the efficacy of the KSHV disease-specific gene therapy vector. We sincerely hope that our vector helps to widen the pharmaceutical window and ease patients suffering from chemotherapeutic drug side effects and devastating KSHV-associated diseases.

## Materials and Methods

### Chemicals, reagents, and antibodies

Dulbecco’s modified minimal essential medium (DMEM), RPMI 1640 medium, fetal bovine serum (FBS), phosphate-buffered saline (PBS), Trypsin-EDTA solution, 100 X penicillin-streptomycin–L-glutamine solution, Alexa 405-conjugated secondary antibody, Alexa 555-conjugated secondary antibody, Alexa 647-conjugated secondary antibody, SlowFade Gold anti-fade reagent, Lipofectamine 2000 reagent, and high-capacity cDNA reverse transcription kit were purchased from Thermo Fisher (Waltham, MA, USA). Puromycin and G418 solution were obtained from InvivoGen (San Diego, CA, USA). Hygromycin B solution was purchased from Enzo Life Science (Farmingdale, NY, USA). Herpes simplex virus type 1/2 thymidine kinase monoclonal antibody was purchased from Invitrogen (San Diego, CA, USA). The Quick-RNA Miniprep kit was purchased from Zymo Research (Irvine, CA, USA), and the QIAamp DNA mini kit was purchased from QIAGEN (Germantown, MD, USA).

### Cell culture

iSLK cells and 293 cells were maintained in DMEM supplemented with 10% FBS, 1% penicillin-streptomycin-L-glutamine solution in 37℃, and 5% CO_2_. iSLK cells were obtained from Dr. Don Ganem (Novartis Institute for Biomedical Research) and cultured under 1 μg/ml puromycin ^80^. iSLK/KSHV and 293/KSHV cells were also maintained in DMEM supplemented with 10% FBS, 1% penicillin-streptomycin-L-glutamine solution under 1 μg/ml puromycin and 500 μg/ml hygromycin. For reactivation, iSLK/KSHV cells were cultured for 4 or 5 days in the presence of 1 μM sodium butyrate and 1 μg/mL doxycycline in DMEM. Insect Sf9 cells were maintained in serum-free PSFM-J1 medium with 50 mL suspension cultures (Fuji Film Wako Chemical). iPSCs were generated from peripheral blood mononuclear cells by introducing the Yamanaka factors (OCT4, SOX2, KLF4, and c-MYC) using episomal vectors. After transduction, the cells were cultured in Stemfit (Ajinomoto, Japan) on iMatrix-511-coated plates under hypoxic conditions (5% O_₂_) at 37°C. Colonies with iPSC-like morphology were manually picked and expanded.

### ECFC differentiation from iPSCs

After 2 days of culture in Stemfit media, iPSCs were stimulated by activin A (10 ng/ml) in the presence of FGF-2, VEGF 165, and BMP4 (10 ng/ml) for 24 h. The following day, the media with cytokines was removed and replaced with Stemline II complete media (Sigma) containing FGF-2, VEGF165, and BMP4 (10 ng/ml) to promote endothelial cell emergence and expansion. Media was replaced with fresh Stemline II complete media on days 3, 5, 7, and 9. On day 12, the media was aspirated, and three parts of EGM-2 and one part of Stemline II complete media were added to the cultures. ECFC colonies appeared as tightly adherent cells.

### Construction of pAAV-TR2-*OriP*-TK plasmid

A luciferase reporter plasmid, which encodes Ori-RNA promoter ^44^, was digested with restriction enzymes, *Nco*I and *Xba*I. The luciferase DNA fragment was replaced with a synthesized TKSR39 DNA fragment (IDTDNA)^81^, which also introduces *Cpo*I restriction enzyme sites at the 5’ and 3’ ends of the coding sequence. The cloning procedure also eliminates XbaI site. The DNA fragment encoding Ori-RNA promoter, TKSR39 coding sequence, and Poly(A) site were amplified with primers listed in Supplemental Data, and cloned into pBlue Script TR2 plasmid at *Kpn*I-*Spe*I site ^18^, which generates pTR2-OriP-TK. The AAV.CMV.PI.EGFP.WPRE.bGH plasmid was a gift from Dr. James M. Wilson (Addgene Plasmid #105530), and the plasmid was digested with restriction enzymes *Nhe*I and *Bam*HI, which removes CMV promoter and EGFP coding sequences. New multiple cloning sequences were prepared by annealing two single-stranded DNA (Supplemental Data), and the resulting double-stranded DNA fragment was introduced to the AAV.CMV.PI.EGFP.WPRE.bGH *Nhe*I-*Bam*HI sites with Gene Assembly. The cloning generated unique sites for BamHI and XbaI in the pAAV.CMV.PI.EGFP.WPRE.bGH, resulting in a pAAV promoter/coding null vector. pTR2-*OriP*-TK was digested with *Xba*I and *Bam*HI and cloned into a pAAV promoter/coding null vector, resulting in pAAV8-TR2-*OriP*-TK. Finally, SV40 poly(A) sites from the pGL3 vector were deleted with restriction enzyme digestion and swapped with WSPR.bGH, resulting in AAV.TR2.OriP.TK.WPRE.bGH. The TK coding sequence, which is cloned into CopI site, was exchanged with other *Cpo*I DNA fragments encoding mCardinal and other reporter genes. To transfer the AAV transfer fragment into a Baculovirus transfer vector, pFAST-BAC1 (Invitrogen), was digested with *Bam*HI and *Hind*III and blunted with T4 DNA polymerase. pAAV.TR2.OriP.TK.WPRE.bGH plasmid was digested with *Pac*I, and the entire AAV transfer fragments including two inverted terminal repeats at both ends, were cloned into the pFAST-BAC vector blunted site. The cloning procedures were summarized in Figure 1C.

### Preparation of recombinant baculoviruses with Bac to BAC system

The pFAST-BAC transfer vector (pSR660) ^82^, which expresses the bicistronic AAV-2 rep gene and polycistronic AAV-8 capsid genes, was a gift from Dr. Robert Kotin (Addgene Plasmid #65216)^82^. pFAST-BAC AAV.TR2.OriP.TK.WPRE.bGH or pSR660 were used to homologous recombination in BAC-to-BAC system in E.coli (invitrogen), and purified baculovirus DNAs from bacteria were transfected Sf9 cells. Recombinant baculoviruses were amplified twice and stored in 4°C for immediately use and at −80°C for stocks. Only passage 3 recombinant baculoviruses were used as working stocks to generate large-scale recombinant AAVs.

### Preparation and purification of recombinant AAVs

Sf9 cells (2.0 x 10^8^) cells were co-infected with recombinant baculoviruses encoding AAV transfer sequence and structure proteins and culture for 3 days in 50 mL suspension. Infected Sf9 cells were collected by centrifuge at 1,800g for 10 min at 4°C. Cell palettes were resuspended with resuspension buffer [50 mM Tris-HCl (pH 8.1), 150 mM NaCl, 2 mM MgCl_2_]. Cells were lysis with four cycles of freeze and thaw, and lysed cells were incubated with Benzonase at 37°C for 1 hour. Cell lysates were centrifuged at 4,500 rpm for 15 min at 4 C. Supernatants were loaded on top of the iodixanol gradient as described previously^83^. The step-wise gradient was centrifuged at 28,000 rpm for 16 hours at 4 °C with a SW28 rotor. The 40% iodixanol gradient was fractionated from bottom to top for every 500 uL, and the presence and purity of recombinant AAV were examined with 10% SDS-PAGE. A representative figure is shown in Figure 3B. Iodixanol fractions that contain AAV were pooled with Amicon Ultra Centrifuge Filter units (Millipore Sigma).

### Quantification of KSHV copy number

Culture supernatant containing viral particles were treated with 1 mM MgCl_2_ and DNase I (12 μg/mL) for 15 min at room temperature. The reaction was stopped by the addition of EDTA to 5 mM followed by heating at 70°C for 15 min. Viral genomic DNA was purified using the QIAamp DNA Mini Kit according to the manufacturer’s protocol and eluted in water. Elution was used for real-time qPCR to determine viral copy number, as described previously ^84^.

### Immunofluorescence Staining

Cells were seeded onto glass coverslips in a 6-well plate. After treatment, the cells were washed with PBS and fixed with 2% paraformaldehyde for 15 minutes at room temperature. The fixed cells were then permeabilized with 0.1% Triton X-100 in PBS for 10 minutes and blocked with 5% bovine serum albumin (BSA) for 1 hour at 37°C. Primary antibodies (1:100 dilution) were diluted in blocking buffer and incubated with the cells at 4°C overnight. After washing three times with PBS, the cells were incubated with fluorophore-conjugated secondary antibodies (1:100 dilution) for 1 hour at room temperature in the dark. Nuclei were counterstained with 4’,6-diamidino-2-phenylindole (DAPI) for 5 minutes at room temperature, followed by two additional washes with PBS. Coverslips were mounted onto glass slides using an anti-fade mounting medium. Fluorescence images were captured using a fluorescence microscope (BZ-X700), and image analysis was performed with BZ-X analyzer software.

### RT-qPCR

iSLK cells 293 cells were infected with or without AAV for 2 days at MOI 10^4^ and 2.0 x 10^5^ cells were reseeded in 6 well plate. GCV, OTX105, and/or SAHA were added on the same day the cells were seeded and continued to culture at 37°C, 5% CO_2_. Total RNA was extracted using the Quick-RNA miniprep kit (Zymo Research, Irvine, CA, USA). A 500 ng of RNA was incubated with DNase I for 15 minutes and reverse transcribed with the High Capacity cDNA Reverse Transcription Kit (Thermo Fisher, Waltham, MA USA). SYBR Green Universal master mix (Bio-Rad) was used for qPCR according to the manufacturer’s instructions. Each sample was normalized to 18S ribosomal RNA. All reactions were run in triplicate. Primer sequences used for qRT-PCR are provided in **Supplemental Data**.

### Flow cytometry

Cells were washed twice with PBS and resuspended in FACS buffer (PBS supplemented with 1% FBS). Flow cytometry was carried out by using a BD Acuri instrument (BD Biosciences) and data analysis was performed using FlowJo v10.8.1 (Tree Star) by gating on live cells based on forward versus side scatter profiles.

### Cryo-electron microscopy (Cryo-EM) imaging

4ul of each sample was applied to a glow-discharged (30mA, 30sec) holey carbon grid (300 mesh Quantifoil R1.2/1.3 copper TEM grid) for plunge freezing in liquid nitrogen using Leica EM GP2 plunger at 18 °C. Cryo-EM images were acquired at 200 kV on a Thermo Scientific Glacios electron microscope equipped with a Gatan K3 direct electron detector. The micrographs were recorded at 56,818x (0.88 Å/pixel) calibrated magnification using K3 with a dose of 40 e/Å^2 and −1.6 um defocus using SerialEM.

### RNA-sequencing

Indexed, stranded mRNA-seq libraries were prepared from total RNA (100 ng) using the KAPA Stranded mRNA-Seq kit (Roche) according to the manufacturer’s standard protocol. Libraries were pooled and multiplex sequenced on an Illumina NovaSeq 6000 System (150-bp, paired-end, >30 × 10^6^ reads per sample).

RNA-Seq data was analyzed using a Salmon-tximport-DESeq2 pipeline. Raw sequence reads (FASTQ format) were mapped to the reference human genome assembly (GRCh38/hg38, GENCODE release 36) and quantified with Salmon ^85^. Gene-level counts were imported with tximport ^86^ and differential expression analysis including Volcano plot were performed with DESeq2 ^87^.

### Xenograft mouse model

All animal studies were conducted according to a UC Davis Institutional Animal Care and Use Committee (IACUC)-approved protocol. NRG (NOD.Cg-Rag1tm1Mom Il2rgtm1Wjl/SzJ Strain #:007799) mouse breeding pairs were purchased from the Jackson Laboratory and the colony was maintained in-house.

8–12-week-old NRG mice were injected subcutaneously with iSLK/KSHV cells, which were infected with or without AAV8-TR2-*OriP*-TK for 4 days. Cells are resuspended in PBS and mixed with the same volume of Matrigel (Corning, #354230). One site (right or left hind leg) of each mouse was implanted 120μl containing 5 × 10^6^ cells. Mice were randomly assigned to PBS control or GCV groups. For the GCV group, each mouse was given 50 mg/kg GCV by intraperitoneal injection twice daily starting on day 2 and continued for 5 days. 50 mg/kg GCV were given twice per week staring from week 2. The tumor volume was measured and calculated as volume (mm3) = L ×W2/2 (L is the largest diameter and W is the smallest diameter of the tumor) every three days. The experiment was terminated 38 days after the tumor cells implant. The mice were euthanized whenever the tumor size was over 20 mm, or the tumor volume was over 2000 mm^3^.

### Statistical analysis

Statistical analyses were performed using GraphPad Prism 9.4.1 software. Results are shown as mean ± SD with dots representing individual measurements. Statistical significance was determined by Student’s t-test, ratio paired t-test, or one-way ANOVA with Tukey’s multiple comparison test, and correction for false discovery rate (FDR) as described in each figure legend. FDR corrected p < 0.05 was considered statistically significant.

## Acknowledgments

We want to thank Mr. Christopher Solver Amaya Bautista and Dr. Fei Guo for technical support. We also thank UC Davis Comprehensive Cancer Center Genomic Shared Resource members for depositing sequence data. This research was supported by public health grants from the National Cancer Institute (CA290700, CA299587), the National Institute of Allergy and Infectious Disease (AI167663), and American Cancer Society grant MBGI-24-1255200-01-MBG Grant DOI #: [doi.org/10.53354/ACS.MBGI-24-1255200-01-MBG.pc.gr.222219] to Y.I. The Genomics Shared Resource is supported by the UC Davis Comprehensive Cancer Center Support Grant (CCSG) awarded by the National Cancer Institute (NCI P30CA093373).

## Author Contributions

T.I., K.N., and Y.I. designed the experiments. T.I. and K.H.W. performed xenograft and flow cytometry analyses. T.I. S.K., R.R.D., A.K., and R.R.D. performed bioinformatics, statistical analyses, and visualization of the transcriptomic datasets. R.R.D. prepared sequencing libraries and performed initial bioinformatics analyses. J.E., S.N., and Y.I. designed and constructed vectors and established protocol for AAV productions. K.H.W., J.E., and Y.I. purified AAVs. T.I. and Y.I. wrote the manuscript and all authors edited the first and subsequent drafts.

## Materials & Correspondence

Correspondence and requests for materials should be addressed to Y.I.

## Data availability

Data supporting the findings of this work are available within the paper and its Supplementary Information files. The datasets generated and analyzed during the current study are available from the corresponding author upon request. The RNA-seq datasets generated in this study have been deposited under the accession number, GSE289947.

**Supplementary Figure 1. Construction of pAAV-TR2-*OriP*-TK vector. a. Schematic diagram of pAAV-TR2-*OriP*-TK vector.** mCardinal sequence encoding in the pAAV-TR2-*OriP*-mCardinal vector was replaced with TK sequence. TR: terminal repeat, TK: thymidine kinase, ITR: inverted terminal repeat. **b. Schematic diagram of TK/GCV system.** TK phosphorylates the prodrug ganciclovir (GCV) into a toxic nucleotide analog, leading to selective cell death in TK-expressing cells. The phosphorylated GCV can also diffuse into neighboring bystander cells by gap junction, inducing cytotoxic effects even in non-TK-expressing cells

**Supplementary Figure 2. SAHA and OTX015 stimulate transcription from the TR2-*OriP* vector. a. Fluorescent and bright field cell images.** KSHV-infected 293 cells were seeded in 12 well plates and transduced with AAV8-TR2-*OriP*-mCardinal. Two days after AAV8-TR2-*OriP*-mCardinal infection, cells were treated with mock (DMSO), OTX015 (200 nM), or SAHA (1 μM). Images were taken four days after the AAV infection. Scales: 200 μm**. b. KSHV-infected 293 cell growth.** KSHV-infected 293 cells were seeded in 6 well plates, and GCV (10 μg/ml) with or without OTX015 (200 nM) or SAHA (1 μM) were added to cells two days after AAV8-TR2-*OriP*-TK infection. Cell growth (upper) and GFP signal intensity (lower) were continuously monitored by Incucyte for 90 hours.

**Supplementary Movie 1. iSLK cells growth**. iSLK/KSHV cells were seeded in 6 well plates and transduced with AAV8-TR2-*OriP*-TK. Cells were treated with (a) mock, (b) GCV (5 μM), (c) GCV and OTX015 (200 nM), or (d) GCV and SAHA (1 μm) for four days. Images were taken every 3 hours continuously by Incucyte.

