## Supplementary figures and images for "Design, development, and evaluation of gene therapeutics specific to KSHV-associated diseases"

### Supplemental Figure 1-2

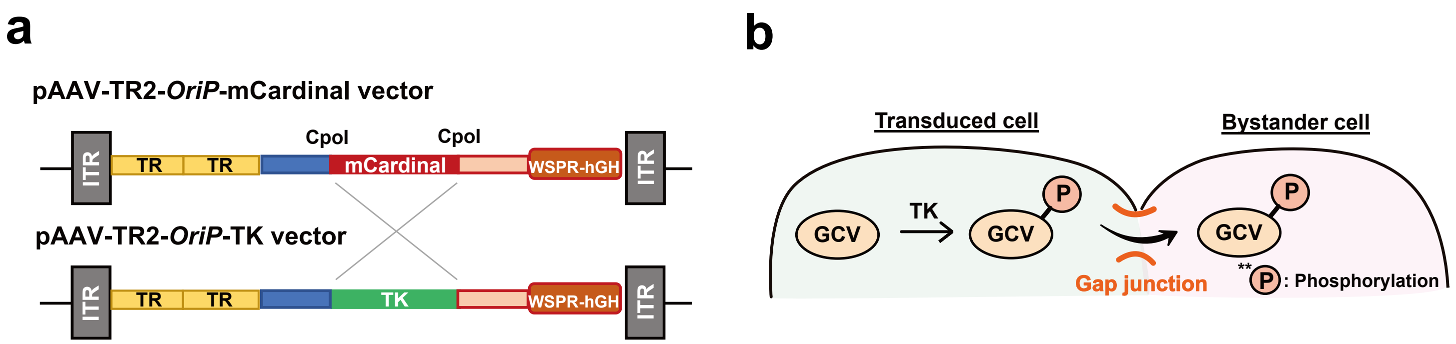


**Supplementary Figure 1**

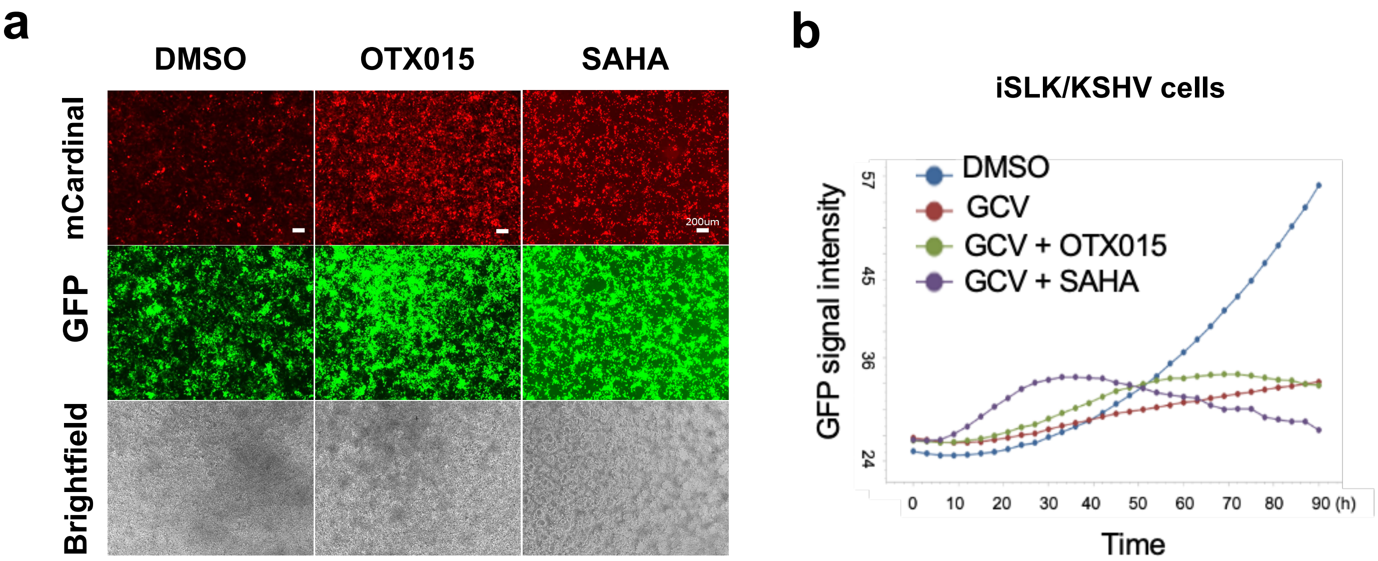


Supplementary Figure 2
